# Economic status mediates the relationship between educational attainment and posttraumatic stress disorder: a multivariable Mendelian randomization study

**DOI:** 10.1101/503300

**Authors:** Renato Polimanti, Andrew Ratanatharathorn, Adam X. Maihofer, Karmel W. Choi, Murray B. Stein, Rajendra A. Morey, Mark W. Logue, Caroline M. Nievergelt, Dan J. Stein, Karestan C. Koenen, Joel Gelernter, the Psychiatric Genomics Consortium Posttraumatic Stress Disorder Workgroup

## Abstract

**Objectives:** To investigate the genetic overlap and causal relationship between posttraumatic stress disorder (PTSD) and traits related to educational attainment.

**Design:** Genetic correlation, polygenic risk scoring, and causal inference via multivariable Mendelian randomization (MR).

**Setting:** Psychiatric Genomics Consortium for PTSD, UK Biobank, 23andMe, and Social Science Genetic Association Consortium.

**Participants:** 23,185 PTSD cases and 151,309 controls; up to 1,131,881 individuals assessed for educational attainment and related traits.

**Main outcome measures:** Genetic correlation obtained from linkage disequilibrium score regression, phenotypic variance explained by polygenic risk scores, and casual effects (beta values) estimated with MR

**Results:** PTSD showed strong negative genetic correlations with educational attainment (EdAtt; r_g_=−0.26, p=4.6×10^−8^). PRS based on genome-wide significant variants associated with EdAtt significantly predicted PTSD (p=6.16×10^−4^), but PRS based on variants associated with PTSD did not predict EdAtt (p>0.05). MR analysis indicated that EdAtt has negative causal effects on PTSD (beta=−0.23, p=0.004). Investigating potential mediators of the EdAtt-PTSD relationship, we observed that propensity for trauma exposure and risk-taking behaviors are risk factors for PTSD independently from EdAtt (beta = 0.36, p = 2.57×10^−5^ and beta = 0.76, p = 6.75×10^−4^, respectively), while income fully mediates the causal effect of EdAtt on PSTD (MR: Income – beta = −0.18, p =0.001; EdAtt – beta =−0.23, p=0.004; multivariable MR: Income – beta = −0.32, p = 0.017; EdAtt – beta = −0.04, p = 0.786).

**Conclusions:** We report novel findings based on large-scale datasets regarding the relationship between educational attainment and PTSD, supporting the role of economic status as the key mediator in the causal relationship observed.

**What is already known on this topic:** There is a well-established negative association of educational attainment and other traits related to cognitive ability with posttraumatic stress disorders (PTSD). However, the findings of these previous studies support various possible causal explanations: 1) individuals with high educational attainment are more resilient with respect to developing PTSD, 2) PTSD negatively impacts cognitive ability, or 3) PTSD and educational attainment share some underlying determinants, including relevant molecular mechanisms.

A key obstacle to disentangling the complex association between educational attainment and PTSD is reverse causation, i.e. the situation in which the outcome precedes and causes the exposure instead of the other way around.

**What this study adds:** We conducted a causal-inference investigation based on large-scale information from the investigation of more than one million individuals. Our main assumption is that genetic information can strongly minimize the bias of reverse causation, because genetic variants are determined at conception and do not change throughout life.

Our findings indicate 1) the effect of traits related to educational attainment on PTSD, 2) no reverse effect of PTSD on educational attainment, and 3) economic status mediates the relationship between educational attainment and PTSD, independently from the brain mechanisms related to educational attainment.

## Introduction

Posttraumatic stress disorder (PTSD) is a psychological condition that occurs in some individuals after exposure to a major traumatic event. Although a large proportion of people are exposed to at least one traumatic event in their lifetimes (estimates from 28% to 90% considering different countries),^1^ only some trauma-exposed individuals will be affected by PTSD. Susceptibility to develop PTSD after trauma exposure is linked to multiple factors, including several pre-trauma risks such as 1) cognitive abilities; 2) coping and response styles; 3) personality factors; 4) psychopathology; 5) psychophysiological factors; and 6) social ecological factors.^2^ Indeed, prospective studies have demonstrated that many variables, previously considered outcomes of trauma, are likely to be pre-trauma risk factors.^2^ Among these cause/consequence conundrums, the relationship of PTSD to cognitive ability and educational attainment is one of the most puzzling.

The current psychiatric definition of PTSD includes multiple cognitive disturbances including concentration difficulties, intrusive recollections of traumatic events, and inability to recall important characteristics of the trauma (DSM-5).^3^ Furthermore, subjective memory and concentration deficits are commonly reported by individuals with PTSD.^4^ However, longitudinal studies investigating various traits related to cognitive function (i.e., intelligence quotient measures, retrieval of memories, negative appraisals about self, cognitive abilities as related to military trainability, extinction learning, and processing speed and memory),^5-7^ have found that lower pre-existing cognitive ability is associated with increased vulnerability for PTSD symptoms.

While there is a robust epidemiologic literature on the negative association between cognitive function and PTSD, the causal mechanisms linking cognitive ability and susceptibility to PTSD remain unclear. Reverse causation (i.e., the situation in which the outcome precedes and causes the exposure instead of the other way around)^8-10^ is one of the key obstacle to disentangling the direction of the mechanisms linking these two phenotypes. This phenomenon can bias the results obtained from traditional observational studies that sometimes attribute causation when all that can be observed directly is an association.^11^

Genetic information can remove the bias of reverse causation from analysis of the PTSD-cognition/education relationship. Indeed, genetic variants are allocated at conception and do not change throughout life. Thus, Mendelian randomization (MR; i.e., a design where causal inference is strengthened by instruments based on genetic information)^12 13^ strongly reduces the possibility of biases induced by reverse causation. Large scale genome-wide association studies (GWAS) have now identified genetic variants that can be used to define reliable genetic instruments that can be applied for causal inference in a MR analysis.^14-18^ The basic principle in MR is that genetic variants that either alter the level of, or mirror the biological effects of, a modifiable environmental exposure that itself alters disease risk should be related to disease risk to the extent predicted by their influence on exposure to the environmental risk factor.^19^

Previous large-scale GWAS conducted by the Social Science Genetic Association Consortium (SSGAC) investigated traits related to educational attainment (e.g., number of years of schooling and most advanced math course ever successfully completed), identifying a number of loci and biological pathways regulating brain mechanisms at the basis of the human cognitive ability.^20-23^ In line with the association between cognitive ability and PTSD and the genetic overlap between educational attainment and cognitive ability, a negative association between PTSD and educational attainment was observed by multiple observational studies.^24 25^ Accordingly, the genetic information from the SSGAC studies can be used to disentangle the causal network linking PTSD, traits related to educational attainment, and the potential mediation of other pre-trauma risk factors such as economic status. Based on these assumptions, we conducted a multivariable MR study using large-scale genome-wide data generated from the investigations of more than one million individuals included in the Psychiatric Genomics Consortium for PTSD (PGC-PTSD), the UK Biobank, 23andMe, and the SSGAC.

## Methods

### Cohorts Investigated

Genome-wide information regarding PTSD was derived from the freeze-2 analysis conducted by the PGC-PTSD working group.^26^ This dataset included 60 ancestrally diverse studies from Europe, Africa and the Americas. In each cohort, lifetime and/or current PTSD status was assessed using various instruments and different versions of the DSM (DSM-III-R, DSM-IV, DSM-5). Genetic information was processed (pre-imputation quality control, imputation, post-imputation quality control, and association analysis) using the same pipeline. A detailed description of the methods applied is available elsewhere.^26^ In our analysis, we focused on the data generated from the analysis of individuals of European descent (23,185 cases; 151,309 controls), because the genome-wide analyses of traits related to educational attainment were conducted only on this ancestry group.

Genome-wide information regarding traits related to educational attainment were derived from the GWAS meta-analysis by the SSGAC,^20^ which investigated educational attainment (EdAtt; i.e., the number of years of schooling that individuals completed) as the primary phenotype in a total of 1,131,881 individuals. In the same SSGAC study,^20^ three additional phenotypes were investigated. Two of these were conducted exclusively among research participants of the personal genomics company 23andMe who answered survey questions about their mathematical background. The first variable, math ability (MathAbility, N = 564,698), was derived from the respondent’s answer to the categorical question “How would you rate your mathematical ability?” The second variable, highest math class (MathClass, N = 430,445), was derived from the answer to a question about the most advanced math course ever successfully completed. The third phenotype investigated, cognitive performance, was assessed in participants (N = 257,828) from the COGENT study and the UK Biobank. A detailed description of the methods applied for phenotypic characterization, quality control of the genetic data, association analysis is available in the main SSGAC GWAS.^20^ For the datasets including participants from 23andMe, we had access only to summary association data of the top 10,000 variants. Accordingly, the datasets derived from 23andme were not used for the reverse MR analysis (i.e., estimating the effect of PTSD on the traits related to educational attainment).

A summary of the datasets and the phenotypes tested and the corresponding information regarding sample size, data available, and included cohorts are reported in Table 1.

**Table 1:**
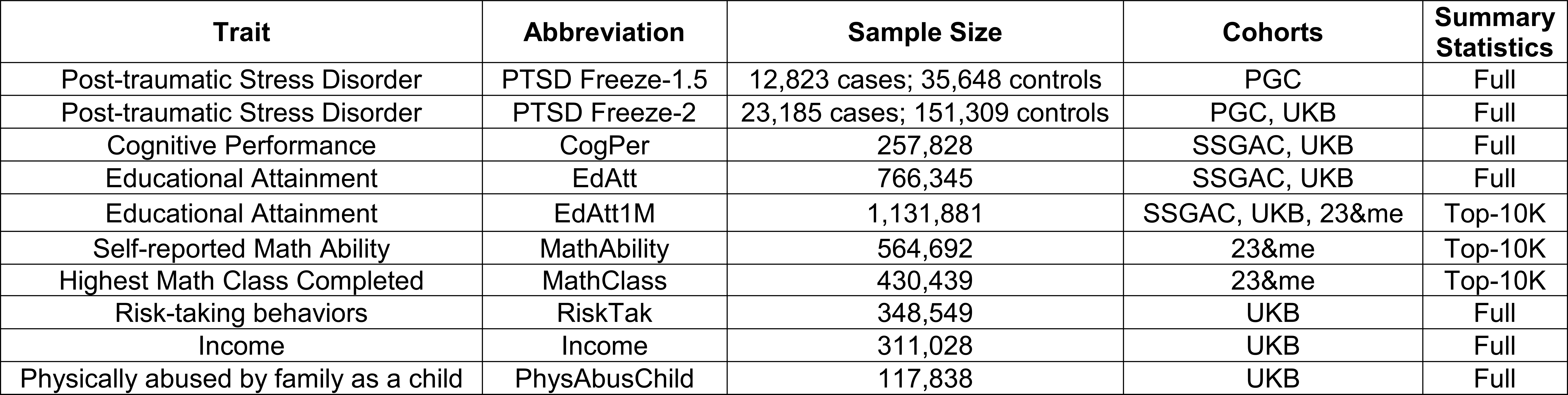
Main traits tested and the corresponding information regarding sample size, data available, and included cohorts (PGC: Psychiatric Genomics Consortium; UKB: UK Biobank; SSGAC: Social Science Genetic Association Consortium).

### Sample Overlap among the Cohorts Investigated

Some of the statistical methods used in the present analysis (i.e., polygenic risk scoring and MR) can be biased by sample overlap between the datasets investigated.^27 28^ The main source of sample overlap between PGC-PTSD and SSGAC GWAS meta-analyses is the presence of some UK Biobank participants in both PGC-PTSD and SSGAC studies. For this reason, some analyses were conducted on a PGC-PTSD subsample excluding the UK Biobank cohort (PGC-PTSD freeze-1.5: 12,823 cases; 35,648 controls). Besides the UK Biobank, the only cohort included in both PGC-PTSD and SSGAC studies is QIMR (Queensland Institute of Medical Research). QIMR cohort represents 1.2% (325 cases and 1,797 controls) of the PGC-PTSD freeze-2 study and 0.7% (N = 8,006) of the SSGAC analysis. We believe that this minor overlap did not affect the results obtained. However, we verified the relationship of PTSD and traits related to educational attainment also considering that some traits were assessed only in the 23andMe cohort, which is independent from PGC-PTSD and UK Biobank samples.

### Genetic Correlation and Definition of the Genetic Instruments

Linkage disequilibrium (LD) score regression was used to estimate the global genetic overlap among the traits investigated. This approach leverages GWAS summary association data together with a LD reference panel to determine pairwise genetic correlations.^29^ For the present study, we included 1,217,311 SNPs in the regression that were present in the HapMap 3 reference panel^30^ and used pre-computed LD scores based on 1000 Genomes Project reference data^31^ on individuals of European ancestry (available at https://github.com/bulik/ldsc). Since sample overlap between GWAS tested does not affect the results obtained from this method,^29^ we were able to calculate pairwise genetic correlations, also including those between datasets including overlapping information from UK Biobank subjects.

To define the genetic instruments for each pairwise comparison to be investigated further via the MR approach, we conducted polygenic risk score (PRS) analyses. The PRSs were calculated after using p-value-informed clumping with an LD cutoff of R^2^ = 0.001 within a 10,000-kb window, excluding the major histocompatibility complex region of the genome because of its complex LD structure, and including only variants with a minor allele frequency >1%. The European samples of the 1000 Genomes Project were used as the LD reference panel. The PRS analysis was conducted on the basis of the GWAS summary association data for both base and target datasets using the gtx R package incorporated in PRSice software.^32^ For each PRS analysis, we calculated an approximate estimate of the explained variance from a multivariate regression model containing the variants included in the PRS.^33^ For the traits related to educational attainment, we considered a genome-wide significance threshold (p < 5×10^−8^). Due to the limited power to detect a large number of genome-wide significant loci, we evaluated multiple P value thresholds (PT = 5×10^−8^, 10^−7^, 10^−6^, 10^−5^, 10^−4^, 0.001, 0.05, 0.1, 0.3, 0.5) to increase the variance explained by the genetic instruments related to the other traits.

### Causal Inference

To assess causality among PTSD, traits related to educational attainment, and potential mediators, we used GWAS summary association data to conduct two-sample MR analyses.^34^ For the variants included in the instrumental variable, we performed LD clumping by excluding alleles that have R^2^ ≥ 0.001 within a 10,000-kb window with another variant with a smaller association P-value. Since different MR methods have different sensitivities to different potential issues, accommodate different scenarios, and vary in their statistical efficiency,^15^ we considered a range of MR methods. The primary analysis was conducted considering a random-effect inverse-variance weighted (IVW) method.^35^ The secondary MR methods included MR Egger,^36^ simple mode,^37^ weighted median,^38^ and weighted mode.^37^ These MR analyses were conducted using the TwoSampleMR R package.^35^ Additionally, due to the fact that some traits showed a limited number of genome-wide significant variants, some of the genetic instruments were based on suggestive significance (GWAS p > 5×10^−8^) calculated on the basis of the PRS analysis. We verified these IVW estimates using the MR-RAPS (Robust Adjusted Profile Score) approach, which is a method designed to identify and estimate confounded causal effects using weak genetic instrumental variables.^39 40^ We conducted multiple sensitivity analyses with respect to the MR tests conducted to exclude the presence of possible biases in the MR estimates. These included the IVW heterogeneity test,^35^ the MR-Egger intercept,^36^ and the MR-RAPS overdispersion test.^39^ For additional confirmation of the absence of distortions due to heterogeneity and pleiotropy, we tested the presence of statistical outliers among the variants included in the genetic instruments using MR-PRESSO (Pleiotropy RESidual Sum and Outlier).^41^ Finally, a leave-one-out analysis was conducted to identify potential outliers among the variants included in the genetic instruments tested. The MR results not biased by horizontal pleiotropy and heterogeneity were entered in the multivariable MR conducted using the method proposed by Burgess and Thompson.^42^ This approach permits evaluation of the independent effects of each risk factor on the outcome, similar to the simultaneous assessment of several treatments in a factorial randomized trial.^42^ In the multivariable MR analysis, the genetic instruments were combined, LD clumping (R^2^ ≥ 0.001 within a 10,000-kb window) was performed to remove correlated variants, and then used to run the multivariable MR analysis using the MendelianRandomization R package.^43^

## Results

### Genetic Correlation and Polygenic Risk Scores between Traits related to Educational Attainment and Posttraumatic Stress Disorder

Our study investigated multiple datasets generated from different cohorts and with different data availability (Table 1). Using LD score regression and datasets with full GWAS summary association data, we observed a negative genetic correlation of PTSD (freeze-2) with EdAtt (rg = −0.26, p = 4.6×10^−8^) and cognitive performance (rg = −0.16, p = 9×10^−4^). The PRS analyses was conducted using PTSD as the target and traits related to educational attainment as the training dataset considering only genome-wide significant (GWS) variants (GWAS p < 5×10^−8^; Figure 1). This analysis was conducted excluding those pairwise comparisons that would have included datasets with the UK Biobank as an overlapping cohort. The strongest PRS association was observed between MathClass GWS PRS with respect to PGC-PTSD freeze-2 (R^2^ = 0.04%, p = 1.13×10^−8^). The same PTSD dataset showed a weaker association with MathAbility PRS (R^2^ = 0.01%, p = 0.004). Significant associations were observed also with respect to PGC-PTSD freeze-1.5 outcome for EdAtt PRS (R^2^ = 0.03%, p = 6.16×10^−4^), MathClass PRS (R^2^ = 0.03%, p = 7.86×10^−4^), and EdAtt1M PRS (R^2^ = 0.03%, p = 0.002). No association of the PRS of cognitive performance and MathAbility was observed with respect to PGC-PTSD freeze-1.5 (R^2^ < 0.01%, p = 0.151; R^2^ < 0.01%, p = 0.105, respectively). We tested the reverse direction (i.e., PTSD as base and traits related to educational attainment as target), but, due to the limited number of GWS loci in the PGC-PTSD analysis, the PRS analysis was conducted considering multiple association PTs to include at least 10 LD-independent variants in each PRS tested (Supplemental Figure 1). We observed a nominally significant association between the PGC-PTSD freeze-1.5 PRS (PT = 10^−5^) and EdAtt (R^2^ = 0.0004%, p=0.041) that would not survive a Bonferroni correction for the number of PTs tested.

**Figure 1:**
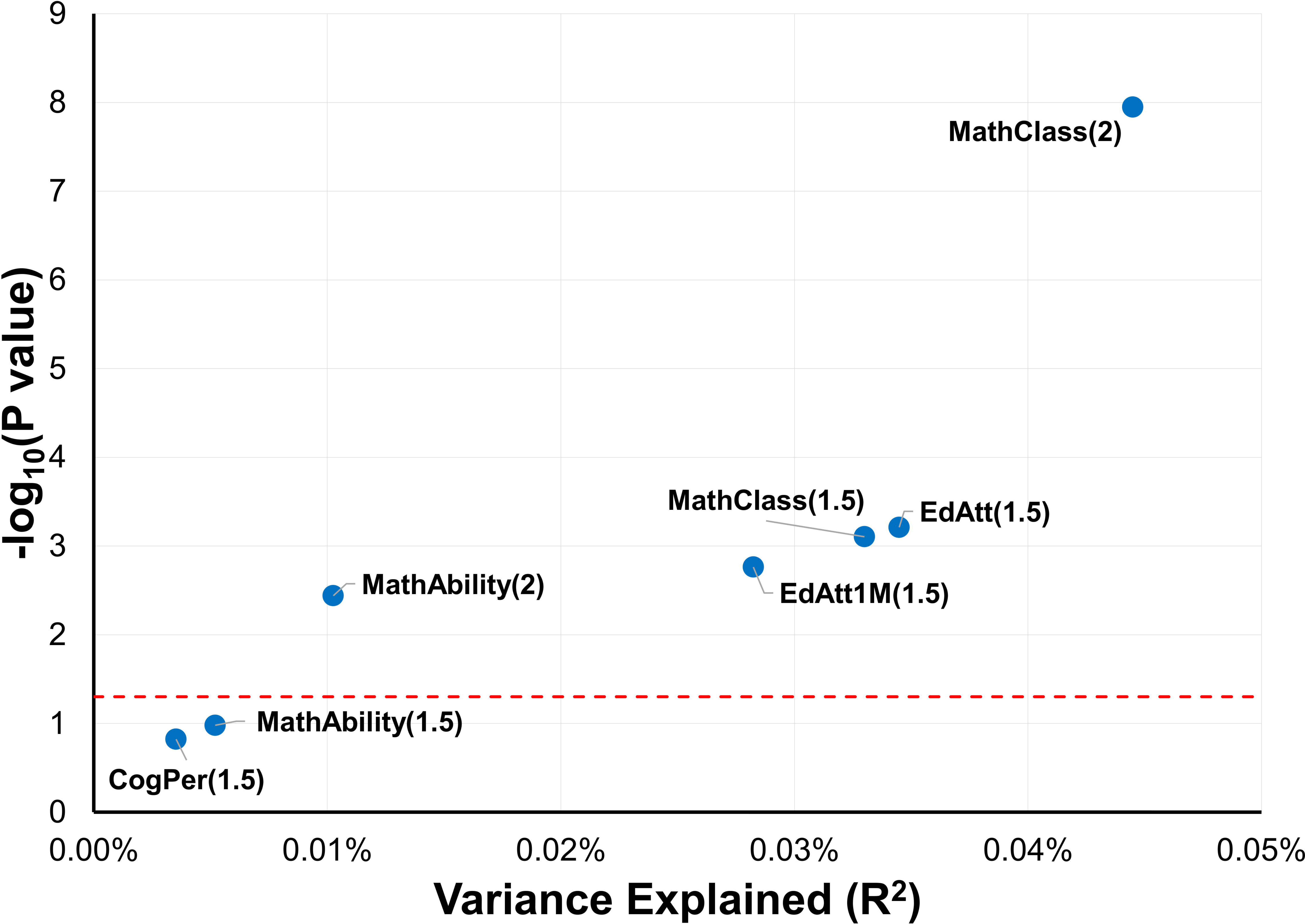
Genetic correlations estimated between traits related to cognitive ability and two versions of the posttraumatic stress disorder dataset, PGC-PTSD freeze-2 (2) and PGC-PTSD freeze-1.5 (1.5). Red dotted line corresponds to nominal significance (p<0.05).

### Causal Inference between Traits related to Educational Attainment and Posttraumatic Stress Disorder

Based on the PRS results, we conducted MR tests using three genetic instruments based on GWS variants. These included MathClass, EdAtt, and EdAtt1M tested with respect to PGC-PTSD datasets (freeze-2 and/or freeze-1.5 depending on UK Biobank overlap). The reverse MR test was conducted on the basis of the PGC-PTSD freeze-1.5 (PT = 10^−5^) with respect to the EdAtt dataset. It was not possible to conduct reverse analyses for the MathClass and EdAtt1M datasets because we did not have access to the full GWAS summary association data for these studies. MathClass showed the strongest causal effect on PTSD (freeze-2; IVW beta = −0.41, p = 3.46×10^−6^). Although we observed a concordant effect when considering other MR methods (Figure 2), the significance was replicated with the MR-RAPS approach (beta = −0.42, p = 3.70×10^−6^), and no outliers were identified by the leave-one-out analysis (Supplemental Figure 2), we observed the presence of possible bias in this result due to heterogeneity and/or pleiotropy (IVW heterogeneity test p = 2.61×10^−4^; MR-RAPS overdispersion test p = 0.007; MR-PRESSO global test p = 5×10^−4^). Although the MR-PRESSO global test identified the presence of potential bias in the analysis, no outlier variants were reported by this method. Thus, we identified the outliers on the basis of the MR-RAPS standardized residuals (−1.96>Z>1.96) and verified their effect on the results of the IVW heterogeneity test (Supplemental Figure 3). Removing the outliers from the MathClass genetic instrument, we confirmed the casual effect of MathClass on PTSD (freeze-2; IVW: beta = −0.39, p = 4.25×10^−7^; MR-RAPS: beta = −0.39, p = 1.06×10^−6^) and the absence of any potential confounders from the sensitivity analyses (Supplemental Table 1; Supplemental Figure 4). The additional MR tests were conducted considering PGC-PTSD freeze-1.5 as outcome and MathClass, EdAtt, and EdAtt1M as exposures. We observed comparable causal effects across the genetic instruments generated from different cohorts (Figure 3; Supplemental Figure 5). However, due to the lower power of PTSD freeze-1.5, we observed a reduction of the significance (MathClass→PTSDfreeze-2: IVW beta = −0.41, p = 3.46×10^−6^; MathClass→PTSDfreeze- 1.5: IVW beta = −0.22, p = 3.79×10^−3^). To further support that MathClass and EdAtt are related to the same causal mechanism, we conducted a multivariable MR analysis, which showed that these two causal effects are not independent from each other (Supplemental Figure 6). Accordingly, we used EdAtt as a proxy of MathClass in the subsequent analyses because we had full access only to the former dataset.

**Figure 2:**
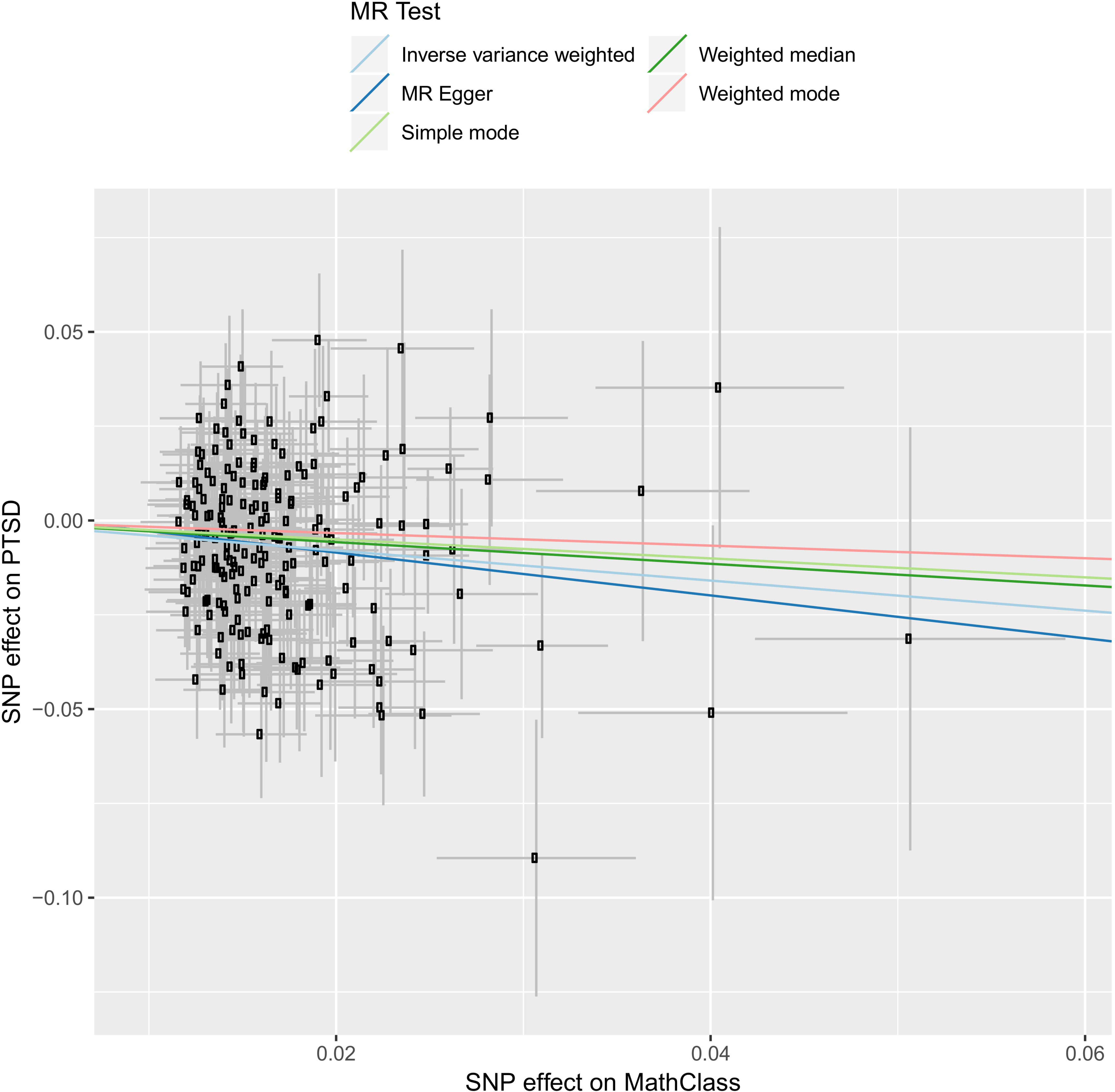
SNP-exposure (MathClass associations, beta) and SNP-outcome (PTSD freeze-2 associations, logOR) coefficients used in the MR analysis. Error bars (95% CIs) are reported for each association.

**Figure 3:**
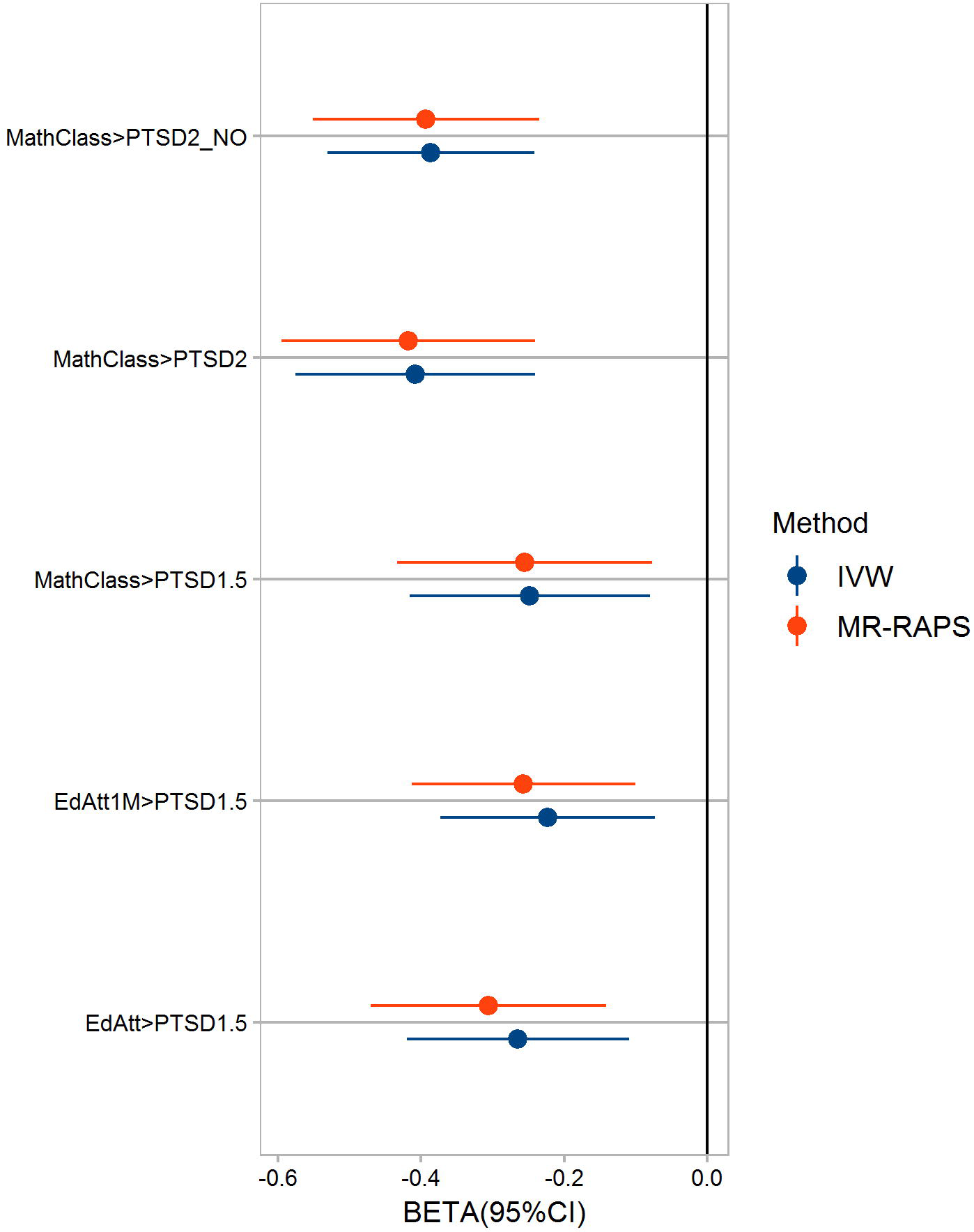
Causal effects estimated using IVW and MR-RAPS methods considering different traits related to educational attainment and two versions of the posttraumatic stress disorder dataset, PGC-PTSD freeze-2 (PTSD2) and PGC-PTSD freeze-1.5 (PTSD1.5).

Palindromic variants with an ambiguous allele frequency (i.e., minor allele frequency close to 50%) can introduce biases in the results generated by two-sample MR analyses.^44^ We did not observe any difference in the causal effects observed using genetic instruments including or excluding these variants (Supplemental Table 2).

Assortative mating has been documented in analyses of genome-wide SNP data on spousal pairs with respect to several traits, including educational attainment.^45^ Since MR methods can be biased by this mechanism,^46^ we confirmed the causal effect of EdAtt on PTSD (freeze-1.5; IVW beta = −0.32, p = 9×10^−4^) after correcting the analysis using the assortative-mating adjustment parameter calculated in the SSGAC GWAS.^20^

We tested the reverse causal mechanism (PTSD→EdAtt), using the best genetic instrument identified by the PRS analysis (freeze-1.5, PT = 10^−5^). No significant causal effect was observed (IVW: beta = 0.006, p = 0.161; MR-RAPS: beta = 0.0006, p = 0.099). To further confirm the absence of reverse causation, we conducted a MR analysis including all LD-independent variants in the genetic instrument and applied the MR-RAPS method only. No causal effect of PTSD on EdAtt was observed (beta = − 0.0006, p = 0.547), but we confirmed the causal effect of EdAtt on PSTD (beta = −0.267, p = 8.06×10^−6^). These outcomes were stable across different adjustments of the MR-RAPS method (Supplemental Table 3).

### Multivariable Mendelian Randomization Analysis and Enrichment Analysis

To investigate the causal effect of EdAtt on PTSD further, we tested three potential mediators (i.e., risk-taking behaviors, income, and trauma exposure) in a multivariable MR analysis. Before entering these potential mediators in the multivariable MR, we verified the reliability of each genetic instrument by conducting a standard two-sample MR. Genome-wide information regarding these traits was derived from GWAS summary association data generated from the UK Biobank (available at http://www.nealelab.is/uk-biobank/). Details regarding the methods used to conduct these analyses are available at https://github.com/Nealelab/UK_Biobank_GWAS. Self-reported risk-taking behaviors were assessed as “Would you describe yourself as someone who takes risks?” (UK Biobank Field ID: 2040). Income was assessed as “Average total household income before tax” (UK Biobank Field ID: 738). Due to the limited number of GWS variants with respect to these traits, we conducted a PRS considering multiple PTs as described previously to define the best genetic instrument for each trait with respect to PGC-PTSD freeze-1.5 dataset (Supplemental Figure 7). The best results were observed for PT = 10^−4^ with risk-taking behaviors (R^2^ = 0.06%, p = 1.53×10^−5^) and PT = 0.001 for income (R^2^ = 0.09%, p = 5.67×10^−8^). The MR analysis based on these genetic instruments confirmed that PTSD is affected by risk-taking behaviors (IVW: beta = 0.755, p = 1.13×10^−4^; MR-RAPS: beta = 0.764, p = 6.75×10^−4^) and income (IVW: beta = −0.18, p = 0.001; MR-RAPS: beta = −0.188, p = 0.003). No evidence of heterogeneity or pleiotropy was observed in either analysis (Supplemental Table 4).

Several traumatic experiences were assessed in the UK Biobank (Supplemental Table 5) and they showed high genetic correlations among each other (Supplemental Table 6). To define the most reliable and the strongest genetic instrument among those related to traumatic experiences, we conducted a PRS analysis, considering four relevant traumatic experiences identified on the basis of their genetic correlation with PTSD, EdAtt, and the other two potential mediators (Supplemental Figure 8). The most informative PRS was observed at PT = 0.001 (Supplemental Figure 9). Then, we conducted a MR analysis testing different trauma-related genetic instruments with respect to PGC-PTSD freeze-1.5. We observed significant causal effects not affected by confounders (Supplemental Table 7) for: “Physically abused by family as a child” (UK Biobank Field ID: 20488; IVW: beta = 0.355, p = 2.99×10^−5^; MR-RAPS: beta = 0.373, p = 6.07×10^−5^) and “Belittlement by partner or ex-partner as an adult” (UK Biobank Field ID: 20521; IVW: beta = 0.141, p = 0.018; MR-RAPS: beta = 0.173, p = 0.005). To verify the presence of a driving signal across these genetic instruments, we conducted a multivariable MR, identifying physical childhood abuse as the strongest genetic instrument for trauma exposure (Supplemental Figure 10).

Having verified the reliability of the genetic instruments related to risk-taking behaviors, income, and childhood trauma, we conducted a multivariable MR analysis with respect to the EdAtt→PTSD causal relationship. We observed that trauma exposure and risk-taking behaviors are independent risk factors for PTSD (i.e., the results obtained from the IVW and multivariable IVW analyses are both significant; Figure 4A, 4B, and 4C). Conversely, income completely mediates the causal effect of EdAtt on PTSD (Figure 4A and 4D). EdAtt has a significant causal effect with respect to PTSD (beta = −0.23, p=0.004), but, when adjusted by income, this result is null (beta = −0.04, p = 0.786). Conversely, income effect on PTSD is still significant when adjusted by EdAtt (unadjusted: beta = −0.18, p = 0.001; adjusted: beta = −0.32, p = 0.017).

**Figure 4:**
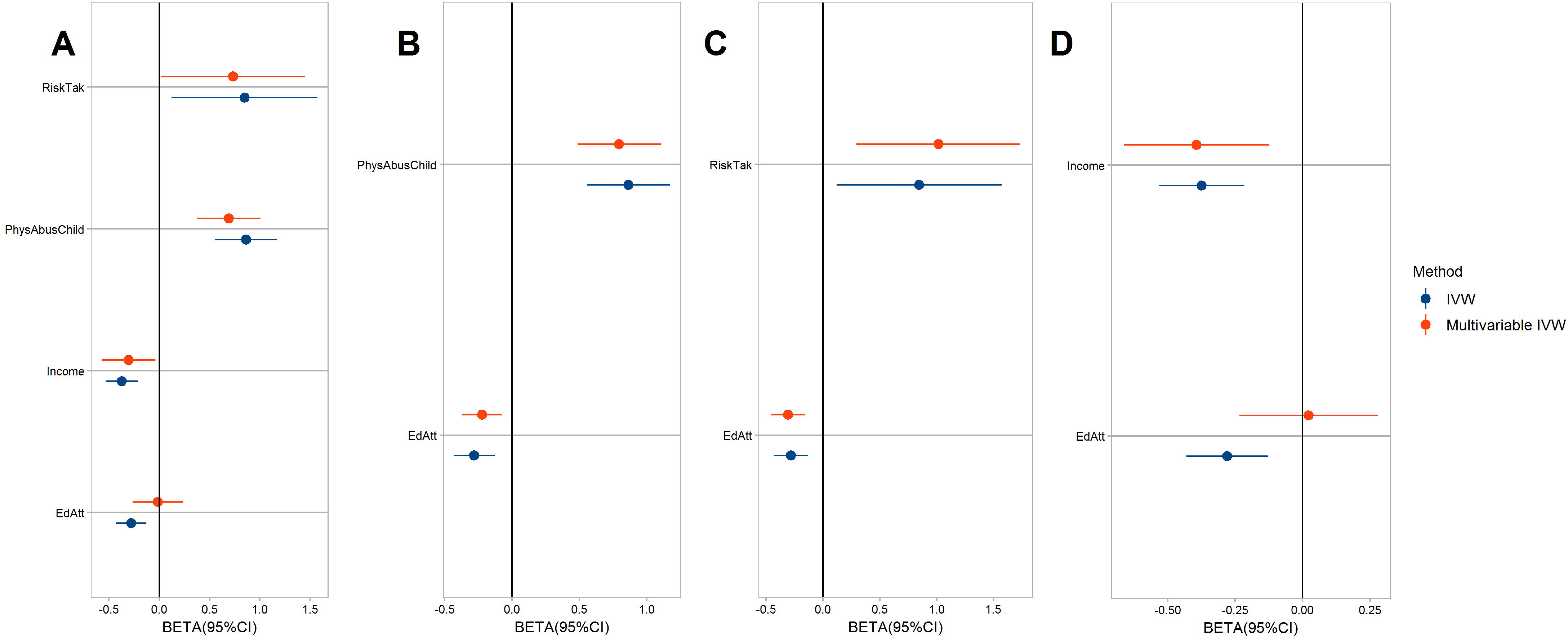
Multivariable Mendelian randomization analysis considering the effects of educational attainment (EdAtt), income, childhood physical abuse (PhysAbusChild), and risk-taking behaviors (RiskTak) on PTSD (**A**); the effects of PhysAbusChild and EdAtt on PTSD (**B**); the effects of RiskTak and EdAtt on PTSD (**C**); and the effects of income and EdAtt on PTSD (**D**).

Although there is a strikingly large genetic correlation between EdAtt and income (rg = 0.81, p < 6.1×10^−308^), income is significantly more correlated with PTSD than EdAtt. (EdAtt: rg = −0.26, p = 4.60×10^−8^; Income: rg = −0.45, p = 9.98×10^−16^; Z_EDATTvsINCOME_ = 2.65, p_EDATTvsINCOME_ = 0.008). We tested if functional differences are present between these two traits conducting enrichment analyses based on tissue and cell-type specific gene expression reference panels^47-51^ and conducted using the MAGMA tool^52^ implemented in FUMA.^53^ Both traits are enriched for brain tissues (e.g., cerebellar hemisphere: EdAtt – beta = 0.098, p = 1.49×10^−16^; income – beta = 0.04, p = 3.83×10^−7^) and neuronal cell-types (e.g., GABAergic neurons: EdAtt – beta = 0.14, p = 8.96×10^−5^; income – beta = 0.049, p = 0.017), but the enrichment signals of the EdAtt GWAS are consistently much stronger than the ones observed in the income GWAS (Figure 5; Supplemental Table 8), clearly showing that EdAtt data is more informative for the brain/cognitive processes than income data. These results suggested that while EdAtt has been shown to have enrichment for brain processes, the fact that income mediates its effect on PTSD – and that income shows a much lower degree of brain-related enrichment – suggests that the EdAtt effect on PTSD is likely due to brain mechanisms related to educational attainment.

**Figure 5:**
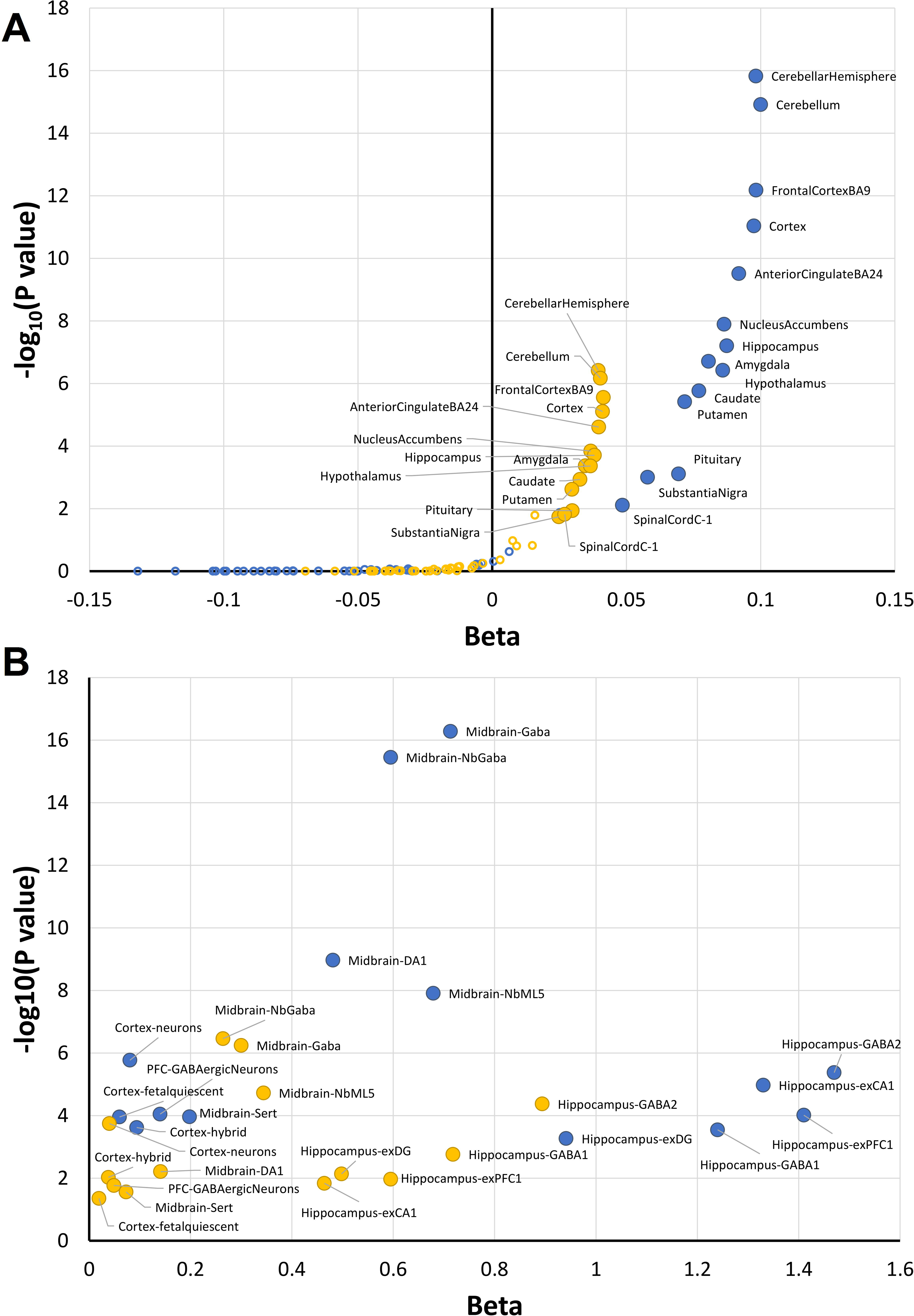
Results of enrichment analyses for educational attainment (blue) and income (yellow). **A**. Tissue-specific enrichments (brain tissues are indicated with full circles; non-Brain tissues are indicated with empty circles). **B**. Cell-type enrichments that were at least nominally significant (p<0.05) in both traits.

## Discussion

The negative association between PTSD and cognitive ability has been confirmed by many independent approaches including cross-sectional and prospective studies. However, the actual causal mechanisms linking these two phenotypes are unclear. Different hypotheses were made: 1) individuals with high educational attainment are more resilient when exposed to traumatic experiences;^24^ 2) PTSD negatively impacts cognitive ability in affected individuals;^54^ 3) PTSD and cognitive ability share some predisposing molecular mechanisms.^55^ The main obstacle to disentangling the biology of the PTSD – cognitive ability association is reverse causation. Genetic information can be deployed to answer this question because it is not affected by reverse causation. We therefore used genome-wide data from more than one million individuals to investigate the causal mechanisms linking PTSD and cognitive ability.

Our analysis was based mainly on data from investigations of traits related to educational attainment (e.g., number of years of schooling and most advanced math course ever successfully completed). These traits are related to numerous biological and non-biological factors.^56^ Previous studies have demonstrated that genetic results deriving from them are mainly informative regarding the brain mechanisms at the basis of the human cognitive ability.^20-23^ Our causation analysis made use of these data to show that traits related to cognitive ability have a causal effect on PTSD. This result was consistent across different cognitive traits investigated in independent cohorts, even when adjusting the analysis for the known effect of assortative mating on these traits. In line with a causal direction from cognition to PTSD, no evidence of any reverse effect was observed.

Although our findings are consistent with a specific direction of causation, we had to evaluate whether other factors could be responsible for this association. Accordingly, we tested three phenotypes of known relevance for PTSD: propensity to trauma exposure,^57^ risk-taking behaviors,^58^ and economic status.^59^ Our MR analysis confirmed the causal effects of each of these risk factors on PTSD, so they were suitable to conduct a multivariable MR analysis of EdAtt with respect to PTSD. This approach permits evaluation of the independent effects of each risk factor on the outcome, similar to the simultaneous assessment of several treatments in a factorial randomized trial.^42^ This analysis clearly showed that, while risk-taking behaviors and propensity to trauma exposure are independent PTSD risk factors, the causal effect of EdAtt on PTSD is completely related to the causal effect of economic status, which is the driving force of the association. EdAtt and income showed a strong genetic overlap, but, while PTSD showed a significantly stronger correlation with income than with EdAtt, our *in silico* investigation showed that EdAtt, but not income, is much more informative for the brain mechanisms that are considered to be responsible for the predisposition to high cognitive ability. Since income appears to be responsible for the EdAtt→PTSD relationship, we hypothesize that this causal mechanism is not related to brain/cognitive ability but rather to other risk factors. A recent genome-wide investigation of social stratification showed that cognition and socio-economic status are correlated with a wide range of factors including personality, psychological traits, mental health, substance use, physical health, reproductive behaviors, and anthropometric traits.^60^ Accordingly, the income effect on PTSD may be related to one or more of these factors, independent of brain mechanisms related to educational attainment.

This study represents the first MR analysis to investigate the causal mechanisms linking cognitive ability to PTSD. It is based on the largest genome-wide datasets for these traits available at this time. The EdAtt→PTSD relationship observed is in line with several prospective studies^2^ and the effect of socioeconomic status on PTSD is also a very well-established risk factor reported in several observational studies.^24 59^ Compared to these previous investigations, the current analyses are based on a much larger population (N > 10^6^) than would ever be feasible for a traditional experimental design (which would include randomized interventions and measurements of outcomes), and without the ethical quandaries that would accompany such randomizations. Also notably, these results are not expected to be affected by reverse causation because of the genetic information used. The findings clearly show a robust income effect that explains the EdAtt→PTSD association, suggesting that brain mechanisms related to cognitive ability are not directly responsible for the association observed. However, although we used state-of-the-art methodology for causal inference analyses and conducted multiple sensitivity analyses, there is still a possibility, though unlikely, that unknown confounders may contribute to the results reported. Indeed, although longitudinal studies and MR analyses are informative for causal inference, the gold standard remains a randomized trial for causal inference. With respect to our hypothesis, a factorial randomized trial would be required to validate our findings, but the design of such an experiment would be very complex, potentially ethically problematic, time consuming, and very expensive.

## Conclusions

This study provides new evidence to elucidate the causal network linking cognitive ability and PTSD, pointing towards risk factors related to economic status rather than brain pathways. We did not observe any effect of PTSD on cognitive ability, but we cannot rule out the possibility that the cognitive processes associated with educational attainment differ from those associated with cognitive decline induced by aging and stress^61 62^ and with more relevant cognitive measures (and we presently do not have a way to assess what exactly those cognitive measures might be), such a relationship could be detected. Further research is necessary to understand the pathways underpinning the link between economic status and PTSD. Additionally, our investigations clearly show that, before claiming causation, MR analysis should consider testing the independence of multiple correlated risk factors with respect to the outcome of interest.

## Acknowledgements

We thank study participants and research groups contributing to PGC-PTSD for sharing their data and the members of the other cited consortia for making their data available.

## Contributors

RP analyzed the data, drew the figures, and wrote the first draft of the manuscript. All authors contributed to the interpretation of the results and critical revision of the manuscript for important intellectual content and approved the final version of the manuscript. For the PGC-PTSD workgroup the investigators contributed to the design and implementation of this specific study and/or provided data but did not participate in analysis or writing of the present study. RP is guarantor.

## Ethical approval

This study is based on publicly available summarized data. Individual studies within each genome-wide association study had received approval from a relevant institutional review board, and informed consent was obtained from participants.

## Data sharing

No additional data available.

## Transparency

The lead author (SCL) affirms that the manuscript is an honest, accurate, and transparent account of the study being reported; that no important aspects of the study have been omitted; that discrepancies from the study as planned have been explained, and that the paper conforms to transparency policy of the International Committee of Medical Journal Editors uniform requirement for manuscripts submitted to biomedical journals.

---

**Supplemental Figure 1**: Effect of the PTSD PRS on educational attainment (yellow) and cognitive performance (green) considering different inclusion thresholds. Red dotted line corresponds to nominal significance (p<0.05).

**Supplemental Figure 2**: Leave-one-out analysis conducted with respect to the MathClass→PTSD2 result.

**Supplemental Figure 3**: Identification of potential outliers (in red) in MathClass genetic instrument based on IVW heterogeneity test and MR-RAPS standardized residuals.

**Supplemental Figure 4**: Results of the MathClass→PTSD2 analysis after the removal of the potential outliers from the genetic instrument. **A**. SNP-exposure (MathClass associations, beta) and SNP-outcome (PTSD freeze-2 associations, logOR) coefficients used in the MR. Error bars (95% CIs) are reported for each association. **B**. Leave-one-out analysis conducted with respect to the result excluding the potential outliers from the genetic instrument.

**Supplemental Figure 5**: Results of the sensitivity analyses with respect to all MR analyses conducted. Red dotted line corresponds to nominal significance (p<0.05).

**Supplemental Figure 6**: Multivariable Mendelian randomization analysis considering the effects of EdAtt and MathClass on PTSD.

**Supplemental Figure 7**: Effect of income and RiskTak PRS (green and blue, respectively) on PTSD considering different inclusion thresholds. Red dotted line corresponds to nominal significance (p<0.05).

**Supplemental Figure 8**: Genetic correlations between traumatic experiences and the other traits of interest.

**Supplemental Figure 9**: Effect of trauma-related PRS on PTSD considering different inclusion thresholds. Red dotted line corresponds to nominal significance (p<0.05).

**Supplemental Figure 10**: Multivariable Mendelian randomization analysis considering the effects of “Physically abused by family as a child” and “Belittlement by partner or ex-partner as an adult” on PTSD.

