## Supplemental Material for "Economic status mediates the relationship between educational attainment and posttraumatic stress disorder: a multivariable Mendelian randomization study"

**Supplemental Table 1**: Results of the sensitivity analyses conducted with respect to the MathClass→PTSD test with and without the outlier variants (Supplemental Figure 3) in the MathClass genetic instrument.

| **MR-Egger** | **Intercept** | **SE** | **P** |
| --- | --- | --- | --- |
| With-Outliers | 0.003 | 0.007 | 0.668 |
| Without-Outliers | 0.003 | 0.006 | 0.608 |
| **IVW Heterogeneity Test** | **Q** | **df** | **P** |
| With-Outliers | 267.4 | 192 | 2.61E-04 |
| Without-Outliers | 160.2 | 174 | 0.765 |
| **MR-RAPS** | **Estimated pleiotropy variance** | **SE** | **P** |
| With-Outliers | 0.00011 | 4.11E-05 | 0.007 |
| Without-Outliers | 0 | 0 | NaN |
| **MR-PRESSO Global test** | **RSSobs** | **P** | **Outliers N** |
| With-Outliers | 281.4 | 5.00E-04 | 0 |
| Without-Outliers | 173.3 | 0.760 | 0 |

**Supplemental Table 2**: Results of the IVW analyses considering genetic instruments with and without palindromic variants with ambiguous allele frequencies (PAL and noPAL, respectively).

| **Test** | **Genetic Instrument** | **Beta** | **SE** | **LCI** | **UCI** |
| --- | --- | --- | --- | --- | --- |
| MathClass>PTSD2 | PAL | -0.41 | 0.09 | -0.58 | -0.24 |
|  | noPAL | -0.40 | 0.09 | -0.57 | -0.23 |
| MathClass>PTSD2_NO | PAL | -0.39 | 0.07 | -0.53 | -0.24 |
|  | noPAL | -0.37 | 0.08 | -0.53 | -0.22 |
| MathClass>PTSD1.5 | PAL | -0.25 | 0.09 | -0.42 | -0.08 |
|  | noPAL | -0.24 | 0.09 | -0.41 | -0.07 |
| EdAtt1M>PTSD1.5 | PAL | -0.22 | 0.08 | -0.37 | -0.07 |
|  | noPAL | -0.21 | 0.08 | -0.37 | -0.06 |
| EdAtt>PTSD1.5 | PAL | -0.26 | 0.08 | -0.42 | -0.11 |
|  | noPAL | -0.23 | 0.08 | -0.39 | -0.07 |

**Supplemental Table 3**: MR-RAPS analysis considering various adjustments based on genome-wide genetic instruments (i.e., all informative LD-independent variants are included in the genetic instrument).

| **Over Dispersion** | **Loss Function** | **Beta** | **SE** | **P** |
| --- | --- | --- | --- | --- |
| EdAtt→PTSD | | | | |
| FALSE | l2 | -0.27 | 0.06 | 1.80E-06 |
| FALSE | huber | -0.27 | 0.06 | 2.61E-06 |
| FALSE | tukey | -0.27 | 0.06 | 2.79E-06 |
| TRUE | l2 | -0.26 | 0.06 | 6.39E-06 |
| TRUE | huber | -0.27 | 0.06 | 2.61E-06 |
| TRUE | tukey | -0.27 | 0.06 | 2.79E-06 |
| PTSD→EdAtt | | | | |
| FALSE | l2 | -0.0013 | 0.0008 | 0.117 |
| FALSE | huber | -0.0011 | 0.0008 | 0.196 |
| FALSE | tukey | -0.001 | 0.0008 | 0.224 |
| TRUE | l2 | -0.0003 | 0.001 | 0.763 |
| TRUE | huber | -0.0005 | 0.001 | 0.660 |
| TRUE | tukey | -0.0005 | 0.001 | 0.648 |

**Supplemental Table 4**: Results of the sensitivity analyses conducted with respect to the Income→PTSD and Risk-Tak→PTSD tests.

| **MR-Egger** | **Intercept** | **SE** | **P** |
| --- | --- | --- | --- |
| Income→PTSD | -0.003 | 0.003 | 0.333 |
| Risk-Tak→PTSD | 0.005 | 0.004 | 0.247 |
| **IVW Heterogeneity Test** | **Q** | **df** | **P** |
| Income→PTSD | 660.4 | 632 | 0.210 |
| Risk-Tak→PTSD | 234.3 | 277 | 0.970 |
| **MR-RAPS** | **Estimated pleiotropy variance** | **SE** | **P** |
| Income→PTSD | 3.32E-05 | 2.76E-05 | 0.333 |
| Risk-Tak→PTSD | 0 | 0 | NaN |
| **MR-PRESSO Global test** | **RSSobs** | **P** | **Outliers N** |
| Income→PTSD | 680.7 | 0.166 | 0 |
| Risk-Tak→PTSD | 240.6 | 0.978 | 0 |

**Supplemental Table 5**: Traumatic experiences assessed in the UK Biobank.

| **Field ID** | **Description** | **N (or case/controls)** |
| --- | --- | --- |
| 20487 | Felt hated by family member as a child | 117,749 |
| 20488 | Physically abused by family as a child | 117,838 |
| 20489 | Felt loved as a child | 117,624 |
| 20490 | Sexually molested as a child | 116,773 |
| 20491 | Someone to take to doctor when needed as a child | 117,301 |
| 20521 | Belittlement by partner or ex-partner as an adult | 117,741 |
| 20522 | Been in a confiding relationship as an adult | 115,099 |
| 20523 | Physical violence by partner or ex-partner as an adult | 117,746 |
| 20524 | Sexual interference by partner or ex-partner without consent as an adult | 117,727 |
| 20525 | Able to pay rent/mortgage as an adult | 116,296 |
| 20526 | Been in serious accident believed to be life-threatening | 11,325/106597 |
| 20527 | Been involved in combat or exposed to war-zone | 4,010/113,944 |
| 20528 | Diagnosed with life-threatening illness | 19,291/98,326 |
| 20529 | Victim of physically violent crime | 21,926/95,920 |
| 20530 | Witnessed sudden violent death | 15,959/101,903 |
| 20531 | Victim of sexual assault | 17,230/99,441 |

**Supplemental Table 6**: Genetic correlation among trauma experiences assessed in the UK Biobank (Supplemental Table 5).

| **Phenotype1-Phenotype2** | **r_g_** | **SE** | **P** |
| --- | --- | --- | --- |
| 20487-20489 | -0.8399 | 0.0316 | 1.59E-155 |
| 20487-20488 | 0.9207 | 0.0387 | 2.43E-125 |
| 20488-20489 | -0.7443 | 0.0355 | 1.24E-97 |
| 20521-20523 | 0.8238 | 0.0491 | 4.58E-63 |
| 20487-20521 | 0.8924 | 0.0557 | 1.14E-57 |
| 20490-20531 | 0.9294 | 0.0645 | 4.22E-47 |
| 20489-20521 | -0.7114 | 0.0497 | 1.91E-46 |
| 20488-20521 | 0.7503 | 0.0589 | 3.38E-37 |
| 20489-20491 | 0.6571 | 0.0534 | 8.92E-35 |
| 20488-20531 | 0.7714 | 0.064 | 1.94E-33 |
| 20523-20524 | 0.9313 | 0.0803 | 4.50E-31 |
| 20489-20523 | -0.733 | 0.0647 | 9.08E-30 |
| 20487-20523 | 0.8611 | 0.0766 | 2.36E-29 |
| 20488-20523 | 0.8143 | 0.0749 | 1.49E-27 |
| 20521-20531 | 0.7104 | 0.0681 | 1.71E-25 |
| 20521-20524 | 0.9425 | 0.0921 | 1.33E-24 |
| 20487-20531 | 0.6732 | 0.0686 | 1.01E-22 |
| 20488-20490 | 0.7403 | 0.0766 | 4.38E-22 |
| 20489-20531 | -0.572 | 0.0597 | 9.49E-22 |
| 20487-20490 | 0.7335 | 0.0852 | 7.60E-18 |
| 20491-20525 | 0.7668 | 0.0915 | 5.29E-17 |
| 20529-20531 | 0.8703 | 0.1047 | 9.39E-17 |
| 20489-20490 | -0.541 | 0.0687 | 3.35E-15 |
| 20523-20531 | 0.7001 | 0.0915 | 1.95E-14 |
| 20490-20521 | 0.646 | 0.0855 | 4.25E-14 |
| 20524-20531 | 0.9057 | 0.1206 | 6.05E-14 |
| 20488-20529 | 0.5572 | 0.075 | 1.11E-13 |
| 20487-20491 | -0.5185 | 0.0707 | 2.24E-13 |
| 20491-20523 | -0.6919 | 0.0954 | 4.13E-13 |
| 20491-20521 | -0.5721 | 0.0805 | 1.16E-12 |
| 20490-20523 | 0.7469 | 0.1066 | 2.42E-12 |
| 20489-20522 | 0.4133 | 0.0595 | 3.71E-12 |
| 20488-20491 | -0.4737 | 0.0704 | 1.74E-11 |
| 20489-20524 | -0.6631 | 0.1015 | 6.33E-11 |
| 20526-20529 | 1 | 0.1706 | 9.55E-11 |
| 20488-20530 | 0.5655 | 0.0878 | 1.16E-10 |
| 20526-20531 | 0.8114 | 0.1282 | 2.49E-10 |
| 20523-20525 | -0.6959 | 0.1108 | 3.42E-10 |
| 20530-20531 | 0.614 | 0.099 | 5.50E-10 |
| 20523-20529 | 0.6458 | 0.1068 | 1.46E-09 |
| 20488-20527 | 0.7115 | 0.118 | 1.64E-09 |
| 20490-20524 | 0.8124 | 0.1356 | 2.11E-09 |
| 20487-20524 | 0.6334 | 0.1062 | 2.47E-09 |
| 20487-20530 | 0.5684 | 0.0959 | 3.08E-09 |
| 20526-20530 | 0.8198 | 0.1411 | 6.26E-09 |
| 20521-20529 | 0.5158 | 0.0896 | 8.73E-09 |
| 20489-20526 | -0.5582 | 0.0978 | 1.13E-08 |
| 20489-20529 | -0.4056 | 0.0711 | 1.17E-08 |
| 20488-20524 | 0.5921 | 0.105 | 1.72E-08 |
| 20529-20530 | 0.6178 | 0.1099 | 1.92E-08 |
| 20521-20526 | 0.6381 | 0.1158 | 3.58E-08 |
| 20487-20529 | 0.4865 | 0.0887 | 4.10E-08 |
| 20491-20522 | 0.4983 | 0.091 | 4.32E-08 |
| 20487-20526 | 0.6625 | 0.1262 | 1.52E-07 |
| 20489-20530 | -0.3474 | 0.0664 | 1.67E-07 |
| 20488-20526 | 0.5618 | 0.111 | 4.13E-07 |
| 20524-20529 | 0.6632 | 0.1361 | 1.10E-06 |
| 20524-20526 | 0.8346 | 0.1762 | 2.18E-06 |
| 20487-20522 | -0.3456 | 0.0736 | 2.62E-06 |
| 20489-20525 | 0.3436 | 0.0736 | 3.06E-06 |
| 20490-20530 | 0.5835 | 0.1262 | 3.76E-06 |
| 20527-20530 | 0.6581 | 0.1428 | 4.06E-06 |
| 20488-20525 | -0.3998 | 0.0884 | 6.15E-06 |
| 20490-20526 | 0.6938 | 0.154 | 6.67E-06 |
| 20521-20530 | 0.4157 | 0.0926 | 7.21E-06 |
| 20487-20527 | 0.5476 | 0.1228 | 8.26E-06 |
| 20523-20526 | 0.6348 | 0.1436 | 9.90E-06 |
| 20523-20530 | 0.4685 | 0.1087 | 1.65E-05 |
| 20526-20527 | 0.9542 | 0.2243 | 2.09E-05 |
| 20527-20531 | 0.5404 | 0.1281 | 2.47E-05 |
| 20491-20531 | -0.3507 | 0.0842 | 3.11E-05 |
| 20524-20530 | 0.5885 | 0.1417 | 3.28E-05 |
| 20526-20528 | 0.853 | 0.2055 | 3.32E-05 |
| 20523-20527 | 0.5835 | 0.1497 | 9.75E-05 |
| 20521-20525 | -0.3805 | 0.1001 | 0.0001 |
| 20487-20525 | -0.3492 | 0.0917 | 0.0001 |
| 20490-20529 | 0.4375 | 0.1129 | 0.0001 |
| 20490-20491 | -0.4118 | 0.1093 | 0.0002 |
| 20521-20528 | 0.4716 | 0.1284 | 0.0002 |
| 20489-20527 | -0.3946 | 0.1048 | 0.0002 |
| 20491-20530 | -0.417 | 0.112 | 0.0002 |
| 20522-20523 | -0.3333 | 0.0915 | 0.0003 |
| 20524-20525 | -0.5356 | 0.147 | 0.0003 |
| 20521-20527 | 0.4853 | 0.1334 | 0.0003 |
| 20527-20529 | 0.5639 | 0.1624 | 0.0005 |
| 20528-20531 | 0.4717 | 0.1352 | 0.0005 |
| 20488-20522 | -0.2382 | 0.0682 | 0.0005 |
| 20488-20528 | 0.43 | 0.1237 | 0.0005 |
| 20522-20525 | 0.3549 | 0.1029 | 0.0006 |
| 20491-20527 | -0.4869 | 0.1415 | 0.0006 |
| 20487-20528 | 0.3784 | 0.1114 | 0.0007 |
| 20523-20528 | 0.4946 | 0.1521 | 0.0011 |
| 20490-20528 | 0.4906 | 0.1498 | 0.0011 |
| 20490-20527 | 0.5086 | 0.1598 | 0.0015 |
| 20528-20530 | 0.497 | 0.1585 | 0.0017 |
| 20489-20528 | -0.3004 | 0.0965 | 0.0019 |
| 20528-20529 | 0.4998 | 0.1642 | 0.0023 |
| 20524-20527 | 0.5686 | 0.1883 | 0.0025 |
| 20525-20530 | -0.3357 | 0.117 | 0.0041 |
| 20491-20524 | -0.3506 | 0.1244 | 0.0048 |
| 20491-20526 | -0.3962 | 0.1423 | 0.0054 |
| 20491-20528 | -0.3578 | 0.1355 | 0.0083 |
| 20524-20528 | 0.487 | 0.1998 | 0.0148 |
| 20521-20522 | -0.1879 | 0.0808 | 0.0201 |
| 20525-20531 | -0.2388 | 0.1037 | 0.0213 |
| 20490-20525 | -0.2303 | 0.117 | 0.0491 |
| 20525-20527 | -0.2874 | 0.1555 | 0.0646 |
| 20527-20528 | 0.307 | 0.179 | 0.0862 |
| 20491-20529 | -0.1828 | 0.1107 | 0.0987 |
| 20522-20524 | -0.1516 | 0.1135 | 0.1816 |
| 20525-20529 | 0.1311 | 0.1135 | 0.2482 |
| 20522-20531 | -0.094 | 0.0864 | 0.2766 |
| 20525-20528 | -0.1683 | 0.1609 | 0.2958 |
| 20522-20529 | 0.1056 | 0.1025 | 0.3028 |
| 20525-20526 | -0.1249 | 0.1605 | 0.4366 |
| 20490-20522 | -0.059 | 0.1023 | 0.5642 |
| 20522-20527 | -0.0608 | 0.1211 | 0.6154 |
| 20522-20530 | 0.0292 | 0.0863 | 0.7352 |
| 20522-20528 | -0.0026 | 0.138 | 0.9848 |
| 20522-20526 | 0.0011 | 0.1264 | 0.9928 |

**Supplemental Table 7**: Results (causal effects and sensitivity analyses) of the MR test conducted using trauma-related genetic instruments with respect to PTSD.

| **Exposure**  **Method** | | | **20487** | | **20488** | | **20489** | | **20521** |
| --- | --- | --- | --- | --- | --- | --- | --- | --- | --- |
| **IVW** | Beta | 0.212 | | 0.356 | | -0.255 | | 0.172 | |
|  | SE | 0.072 | | 0.085 | | 0.052 | | 0.059 | |
|  | Pl | 0.003 | | 2.57E-05 | | 7.06E-07 | | 0.003 | |
|  | Q | 449.0 | | 453.4 | | 461.0 | | 415.2 | |
|  | df | 425 | | 410 | | 458 | | 445 | |
|  | P | 0.203 | | 0.068 | | 0.452 | | 0.841 | |
| **MR-Egger** | BETA | 0.493 | | 0.033 | | 0.110 | | 0.057 | |
|  | SE | 0.151 | | 0.191 | | 0.122 | | 0.124 | |
|  | Pl | 0.001 | | 0.863 | | 0.366 | | 0.643 | |
|  | Intercept | -0.007 | | 0.007 | | -0.010 | | 0.003 | |
|  | SE | 0.003 | | 0.003 | | 0.003 | | 0.003 | |
|  | P | 0.035 | | 0.060 | | 0.001 | | 0.291 | |
| **MR-RAPS** | Beta | 0.238 | | 0.373 | | -0.281 | | 0.173 | |
|  | SE | 0.080 | | 0.093 | | 0.057 | | 0.062 | |
|  | P | 0.003 | | 6.07E-05 | | 8.78E-07 | | 0.005 | |
|  | Variance | 6.65E-05 | | 1.03E-04 | | 2.45E-05 | | 0 | |
|  | SE | 5.62E-05 | | 5.65E-05 | | 4.05E-05 | | 0 | |
|  | P | 0.236 | | 0.069 | | 0.546 | | NaN | |
| **MR-PRESSO** | Beta | 0.205 | | 0.355 | | -0.260 | | 0.141 | |
|  | SE | 0.072 | | 0.084 | | 0.051 | | 0.054 | |
|  | P | 0.005 | | 2.99E-05 | | 5.35E-07 | | 0.010 | |
|  | RSSobs | 452.6324 | | 462.4988 | | 466.4951 | | 428.5089 | |
|  | P | 0.225 | | 0.066 | | 0.470 | | 0.778 | |

**Supplemental Table 8**: Results of the enrichment analysis based on tissue- and cell-type specific transcriptomic data.

| **Trait** | **EdAtt** | | **Income** | |
| --- | --- | --- | --- | --- |
| **GTEx** | | | | |
| **Tissue** | **BETA** | **P** | **BETA** | **P** |
| Brain_Cerebellar_Hemisphere | 0.0982 | 1.49E-16 | 0.0395 | 3.83E-07 |
| Brain_Cerebellum | 0.1 | 1.21E-15 | 0.0403 | 6.76E-07 |
| Brain_Frontal_Cortex_BA9 | 0.0983 | 6.62E-13 | 0.0414 | 2.78E-06 |
| Brain_Cortex | 0.0975 | 9.20E-12 | 0.0411 | 7.88E-06 |
| Brain_Anterior_cingulate_cortex_BA24 | 0.0919 | 3.07E-10 | 0.0397 | 2.47E-05 |
| Brain_Nucleus_accumbens_basal_ganglia | 0.0864 | 1.27E-08 | 0.0367 | 0.000145 |
| Brain_Hippocampus | 0.0874 | 6.24E-08 | 0.0381 | 0.000197 |
| Brain_Amygdala | 0.0806 | 1.96E-07 | 0.0347 | 0.000429 |
| Brain_Hypothalamus | 0.0859 | 3.80E-07 | 0.0366 | 0.000434 |
| Brain_Caudate_basal_ganglia | 0.077 | 1.71E-06 | 0.0327 | 0.001168 |
| Brain_Putamen_basal_ganglia | 0.0717 | 3.81E-06 | 0.0297 | 0.002422 |
| Pituitary | 0.0694 | 0.000776 | 0.0298 | 0.011664 |
| Brain_Substantia_nigra | 0.0579 | 0.000987 | 0.0248 | 0.0183 |
| Brain_Spinal_cord_cervical_c-1 | 0.0485 | 0.007807 | 0.027 | 0.015214 |
| Testis | 0.0251 | 0.013298 | 0.0159 | 0.01622 |
| Cells_EBV-transformed_lymphocytes | 0.00632 | 0.23595 | 0.00765 | 0.10632 |
| Cells_Transformed_fibroblasts | 0.000486 | 0.48597 | 0.00913 | 0.15612 |
| Adrenal_Gland | -0.00383 | 0.56375 | -0.0173 | 0.88267 |
| Ovary | -0.00584 | 0.59539 | 0.015 | 0.15093 |
| Muscle_Skeletal | -0.00493 | 0.64895 | -0.00755 | 0.80473 |
| Uterus | -0.0313 | 0.8422 | 0.00294 | 0.43525 |
| Colon_Sigmoid | -0.0381 | 0.84845 | -0.00345 | 0.56596 |
| Esophagus_Gastroesophageal_Junction | -0.0475 | 0.88722 | -0.0121 | 0.70874 |
| Esophagus_Muscularis | -0.0451 | 0.88997 | -0.0127 | 0.72949 |
| Nerve_Tibial | -0.0358 | 0.89334 | -0.0173 | 0.85216 |
| Heart_Atrial_Appendage | -0.0317 | 0.93275 | -0.0399 | 0.99853 |
| Artery_Tibial | -0.043 | 0.94571 | -0.00671 | 0.66491 |
| Heart_Left_Ventricle | -0.0305 | 0.95833 | -0.0378 | 0.99931 |
| Pancreas | -0.03 | 0.96553 | -0.0162 | 0.92949 |
| Cervix_Ectocervix | -0.0742 | 0.96915 | -0.0122 | 0.70887 |
| Whole_Blood | -0.0205 | 0.98382 | -0.0131 | 0.97082 |
| Thyroid | -0.055 | 0.98426 | -0.00688 | 0.67599 |
| Cervix_Endocervix | -0.0765 | 0.98785 | -0.0154 | 0.78747 |
| Fallopian_Tube | -0.081 | 0.98912 | -0.0217 | 0.86041 |
| Artery_Aorta | -0.0647 | 0.99042 | -0.0149 | 0.82158 |
| Artery_Coronary | -0.0861 | 0.99241 | -0.035 | 0.96024 |
| Liver | -0.0298 | 0.99505 | -0.0247 | 0.99895 |
| Bladder | -0.0888 | 0.9951 | -0.0452 | 0.98972 |
| Vagina | -0.083 | 0.99564 | -0.0341 | 0.96842 |
| Prostate | -0.0991 | 0.99698 | -0.0385 | 0.97004 |
| Skin_Not_Sun_Exposed_Suprapubic | -0.0501 | 0.99786 | -0.0232 | 0.98138 |
| Skin_Sun_Exposed_Lower_leg | -0.0516 | 0.99844 | -0.0232 | 0.98177 |
| Stomach | -0.0949 | 0.99867 | -0.0441 | 0.99077 |
| Adipose_Subcutaneous | -0.0926 | 0.99913 | -0.0454 | 0.99592 |
| Esophagus_Mucosa | -0.0532 | 0.99939 | -0.0296 | 0.99724 |
| Spleen | -0.0513 | 0.99943 | -0.0207 | 0.97981 |
| Breast_Mammary_Tissue | -0.132 | 0.99964 | -0.0696 | 0.99924 |
| Colon_Transverse | -0.103 | 0.99968 | -0.0397 | 0.9879 |
| Kidney_Cortex | -0.074 | 0.99981 | -0.044 | 0.99962 |
| Small_Intestine_Terminal_Ileum | -0.0804 | 0.99985 | -0.0287 | 0.98337 |
| Adipose_Visceral_Omentum | -0.118 | 0.99993 | -0.0586 | 0.99941 |
| Minor_Salivary_Gland | -0.0999 | 0.99998 | -0.0513 | 0.99976 |
| Lung | -0.104 | 1 | -0.0448 | 0.99926 |
| **Cortex** | | | | |
| **Cell type** | **BETA** | **P** | **BETA** | **P** |
| neurons | 0.0804 | 1.70E-06 | 0.0399 | 0.00018 |
| fetal_quiescent | 0.06 | 0.000111 | 0.0192 | 0.044132 |
| hybrid | 0.0938 | 0.00024 | 0.0378 | 0.009398 |
| oligodendrocytes | -0.00061 | 0.51667 | 0.00566 | 0.287 |
| OPC | -0.0114 | 0.75919 | -0.0158 | 0.92398 |
| fetal_replicating | -0.0138 | 0.81459 | -0.00229 | 0.58443 |
| astrocytes | -0.0161 | 0.8728 | -0.00731 | 0.77638 |
| microglia | -0.0474 | 0.99994 | -0.0235 | 0.99536 |
| endothelial | -0.0753 | 1 | -0.0269 | 0.9962 |
| **Hippocampus** | | | | |
| **Cell type** | **BETA** | **P** | **BETA** | **P** |
| GABA2 | 1.47 | 4.23E-06 | 0.894 | 4.24E-05 |
| exCA1 | 1.33 | 1.07E-05 | 0.464 | 0.014588 |
| exPFC1 | 1.41 | 9.71E-05 | 0.595 | 0.010831 |
| GABA1 | 1.24 | 0.000285 | 0.718 | 0.001726 |
| exDG | 0.941 | 0.000542 | 0.498 | 0.007281 |
| exPFC2 | 0.836 | 0.00143 | 0.301 | 0.052282 |
| exCA3 | 0.618 | 0.019504 | 0.0964 | 0.32455 |
| OPC | -0.333 | 0.78642 | -0.0024 | 0.50313 |
| ODC1 | -0.41 | 0.9072 | -0.133 | 0.73382 |
| ODC2 | -0.496 | 0.9702 | -0.0715 | 0.64686 |
| ASC1 | -0.907 | 0.99836 | -0.563 | 0.9925 |
| MG | -1.03 | 0.99839 | -0.625 | 0.98971 |
| NSC | -0.393 | 0.99867 | -0.227 | 0.98834 |
| ASC2 | -0.896 | 0.99968 | -0.681 | 0.99952 |
| END | -0.642 | 1 | -0.182 | 0.96368 |
| **Pre-frontal Cortex** | | | | |
| **Cell type** | **BETA** | **P** | **BETA** | **P** |
| GABAergic_neurons | 0.14 | 8.96E-05 | 0.0488 | 0.017151 |
| Astrocytes | 0.0161 | 0.2512 | -0.00241 | 0.56252 |
| Neurons | 0.00216 | 0.47808 | 0.00402 | 0.43326 |
| OPC | -0.0116 | 0.60234 | -0.00836 | 0.62414 |
| Stem_cells | -0.0404 | 0.93612 | 0.00623 | 0.3562 |
| Microglia | -0.0275 | 0.95191 | -0.0147 | 0.9053 |
| **Midbrain** | | | | |
| **Cell type** | **BETA** | **P** | **BETA** | **P** |
| Gaba | 0.713 | 5.23E-17 | 0.3 | 5.76E-07 |
| NbGaba | 0.595 | 3.56E-16 | 0.264 | 3.46E-07 |
| DA1 | 0.481 | 1.08E-09 | 0.141 | 0.006199 |
| NbML5 | 0.679 | 1.24E-08 | 0.344 | 1.89E-05 |
| Sert | 0.198 | 0.00011 | 0.0729 | 0.02764 |
| DA0 | 0.376 | 0.001459 | 0.0801 | 0.17967 |
| RN | 0.311 | 0.002557 | 0.156 | 0.021646 |
| DA2 | 0.208 | 0.011987 | 0.0173 | 0.39402 |
| NbM | 0.277 | 0.026398 | 0.136 | 0.077594 |
| NbML1 | 0.233 | 0.042149 | -0.00043 | 0.50192 |
| OMTN | 0.042 | 0.36092 | -0.101 | 0.90238 |
| Rgl2b | -0.016 | 0.55399 | 0.0352 | 0.33043 |
| Rgl1 | -0.0731 | 0.7576 | 0.0589 | 0.20091 |
| OPC | -0.083 | 0.8201 | -0.0327 | 0.69907 |
| Rgl2c | -0.12 | 0.91998 | -0.0498 | 0.80025 |
| Rgl2a | -0.163 | 0.96156 | -0.0267 | 0.66654 |
| Rgl3 | -0.142 | 0.9744 | -0.0617 | 0.89154 |
| NProg | -0.366 | 0.99535 | -0.122 | 0.9029 |
| ProgFPL | -0.227 | 0.99689 | -0.0527 | 0.82512 |
| Mgl | -0.168 | 0.99854 | -0.0849 | 0.9805 |
| ProgBP | -0.472 | 0.99997 | -0.146 | 0.96734 |
| ProgM | -0.474 | 0.99999 | -0.131 | 0.96222 |
| Endo | -0.28 | 1 | -0.0964 | 0.99527 |
| Peric | -0.3 | 1 | -0.116 | 0.99723 |
| ProgFPM | -0.429 | 1 | -0.226 | 0.99982 |
